# Dual inhibition of glutaminolysis and autophagy suppresses proliferation of Rheumatoid arthritis fibroblast-like synoviocytes and mitigates arthritis in SKG mice

**DOI:** 10.1101/2025.06.25.661649

**Authors:** Ikuko Naka, Sho Sendo, Takaichi Okano, Kenichi Uto, Soshi Takahashi, Shinya Hayashi, Ryosuke Kuroda, Akio Morinobu, Jun Saegusa

## Abstract

Recent evidence highlights the critical role of immune metabolism in the pathogenesis of rheumatoid arthritis (RA). We previously demonstrated that glutaminase 1 (GLS1), a key enzyme in glutamine metabolism, is upregulated in fibroblast-like synoviocytes from RA patients (RA-FLS) and that GLS1 inhibition exerts an antiproliferative effect on RA-FLS. Glutaminolysis has also been shown to suppress autophagy, raising the possibilitiy that inhibiting glutaminolysis may enhance autophagy. Given this interplay, we hypothesized that dual inhibition of glutaminolysis and autophagy could synergistically suppress RA-FLS proliferation.

This study aimed to investigate the effects of combining autophagy and glutaminolysis inhibition on RA-FLS and arthritis in SKG mice. We utilized chloroquine (CQ) as an autophagy inhibitor and compound 968 (C968), a GLS1 inhibitor, to suppress glutaminolysis. Treatment with C968 upregulated LC3B and ATG5 expression and increased LC3-II protein levels in RA-FLS, indicative of enhanced autophagy. Furthermore, C968 promoted autophagosome formation in RA-FLS. These findings confirm that glutaminolysis inhibition enhances autophagy in RA-FLS.

The combination of C968 and CQ significantly inhibited RA-FLS proliferation and increased apoptotic cell death. Moreover, C968-CQ co-treatment markedly alleviated arthritis severity in SKG mice. Our findings suggest that concurrent suppression of glutaminolysis and autophagy represents a promising therapeutic strategy for RA.

## Introduction

Rheumatoid arthritis (RA) is a chronic systemic autoimmune disease characterized by persistent joint inflammation, immune cell infiltration, and synovial hyperplasia, ultimately leading to cartilage and bone destruction [1]. A key pathological hallmark of RA is synovial hyperplasia and pannus formation, which drive the progression of destructive joint inflammation [2,3]. Among the key players in this process, fibroblast-like synoviocytes (FLS) are the predominant cellular component of the pannus and and exhibit tumor-like aggressive properties, including impaired contact inhibition, enhanced migratory capacity, resistance to apoptosis, and invasiveness to periarticular tissues [4]. Recent studies have identified distinct subsets of RA-FLS, suggesting that targeting RA-specific pathogenic subsets could offer a promising therapeutic strategy for refractory RA [5].

Despite advancements in RA treatment, significant unmet needs remain, largely due to the disease’s multifactorial and heterogeneous nature [6]. Consequently, the development of novel and more effective therapeutic strategies is urgently required.

The antimalarial drug chloroquine (CQ) and its derivative hydroxychloroquine (HCQ)—which share similar pharmacokinetics and mechanisms of action—have long been used in the treatment of RA and other autoimmune diseases [7]. However, CQ or HCQ alone has shown only limited efficacy in RA, necessitating combination therapy for improved treatment outcomes [8].

In recent years, increasing attention has been directed toward the anti-autophagic effects of CQ and HCQ. Autophagy is a fundamental catabolic process in which cells sequester targeted cytoplasmic components within autophagic vacuoles or autophagosomes, degrade them via fusion with lysosomes, and recycle their constituents [9,10]. Several studies have indicated that autophagy plays a cytoprotective role in cancer therapy and various autoimmune diseases [11]. In certain types of cancer, chemotherapy-induced autophagy contributes to drug resistance and autophagy inhibition via CQ or HCQ has been shown to enhance anticancer therapeutic efficacy [12].

Recent studies have also highlighted the critical role of autophagy in RA pathogenesis [13]. Autophagy has been linked to the apoptosis-resistant phenotype of RA-FLS [14,15], and elevated autophagy levels have been observed in synovial tissues of patients with active RA, correlating with disease severity [16]. Additionally, autophagy activation in the synovium has been associated with methotrexate resistance, suggesting its role in treatment refractoriness [17].

Beyond autophagy, cellular metabolism has emerged as a key contributor to RA pathogenesis, including RA-FLS activation [18]. We previously demonstrated that glutaminase 1 (GLS1), a key enzyme in glutamine metabolism, was upregulated in RA-FLS. Furthermore, GLS1 inhibition restrained RA-FLS proliferation and ameliorated inflammatory arthritis in SKG mice [19].

An interplay between autophagy and glutamine metabolism has been suggested, particularly in cancer research. Glutaminolysis inhibits autophagy by activating mTORC1 or counteracting ROS production in cancer cells [20]. Given that glutaminolysis suppression may promote autophagy, we hypothesized that dual inhibition of glutaminolysis and autophagy could exert a synergistic inhibitory effect on RA-FLS proliferation. However, the relationship between autophagy and glutaminolysis in RA-FLS remains unclear.

In this study, we investigated the therapeutic potential of combining autophagy and glutaminolysis inhibition in RA-FLS and a murine RA model. Our findings reveal that the GLS1 inhibitor, compound 968 (C968), enhances autophagy in RA-FLS and that, in combination with the autophagy inhibitor CQ, C968 synergistically suppresses RA-FLS proliferation and enhances apoptosis. Furthermore, we demonstrate that combined C968 and CQ treatment significantly alleviates arthritis in SKG mice, suggesting a promising novel therapeutic strategy for RA.

## Materials and Methods

### Isolation and culture of fibroblast-like synoviocytes

This study was approved by the Ethics Committee of the Graduate School of Medicine, Kobe University (Approval Number: B190157).

Synovial tissues were obtained from adult RA patients undergoing synovectomy or arthroplasty at Kobe University Hospital between September 18, 2020, and March 31, 2021. Oral informed consent was obtained from all participants prior to sample collection. The informed consent process took place between the attending physician and the patient only, without an independent witness. Written consent was deemed unnecessary as the study utilized discarded surgical samples. Patients were informed of the study through public disclosure, and an opt-out option was provided.

The study adhered to the Declaration of Helsinki and the Ethical Guidelines for Medical and Health Research Involving Human Subjects in Japan.

FLS were isolated as previously described [21]. FLS were grown in Dulbecco’s modified Eagle’s medium with 10% fetal bovine serum (Gibco BRL, CA, USA), 2 mM l-glutamine (Gibco BRL) and 1% penicillin-streptomycin (Lonza, Walkersville, MD, USA) at 37 °C in a humidified 5% CO2 incubator. FLS cultures (passages 3-6) were maintained as described previously [21]. FLS were pre-incubated with 10 ng/mL platelet-derived growth factor (PDGF; PeproTech, NJ, USA) for 2 hours prior to exposure to CQ or C968.

### Quantitative real-time PCR

Total RNA was extracted from FLS using the RNeasy kit (Qiagen, Hilden, Germany), and reverse transcription was performed using the QuantiTect Reverse Transcription Kit (Qiagen). Quantitative real-time PCR was carried out using the QuantiTect SYBR Green PCR Kit (Qiagen) on an ABI Prism 9900 instrument (Applied Biosystems, CA, USA), following the manufacturer’s instructions. Gene expression levels were normalized to GAPDH (QT01192646; Qiagen). The primers used were obtained from Qiagen and included:

・ LC3B: 5′-GTCCGACTTATTCGAGAGCAG-3′ (forward) and 5′-CTGAGATTGGTGTGGAGACG-3′ (reverse)

・ ATG5: 5′-GGCCATCAATCGGAAACTCAT-3′ (forward) and 5′-AGCCACAGGACGAAACAGCTT -3′ (reverse).

### Western blotting

Cell lysates from RA-FLS were subjected to western blot analysis using anti-LC3 (MBL, Nagoya, Japan) and anti-β-actin (Sigma-Aldrich) antibodies. Protein-antibody complexes were visualized using the SuperSignal West Dura Extended Duration Substrate (Thermo Fisher Scientific, Waltham, MA, USA) according to the manufacturer’s instructions. Immunoreactive bands were detected using the Amersham Imager 600 (GE Healthcare, Chicago, IL, USA).

### Confocal immunofluorescence

FLS were seeded at 2.5 × 10^4^ cells per well in eight-well chamber slides (Lab-Tek®, Electron Microscopy Sciences, Hatfield, PA, USA). Following treatment with C968 or CQ, cells were washed twice, stained with Cyto-ID® green dye and Hoechst 33342 nuclear dye (Enzo Life Sciences, Farmingdale, USA) for 30 minutes at 37 °C, and subsequently fixed with 4% paraformaldehyde for 20 minutes. After three washes, fluorescence imaging was performed using a Keyence BZ-X700 fluorescence microscope (Keyence, Osaka, Japan).

### Cell proliferation assay

FLS (1 × 10^4^ cells/well) were seeded in 96-well plates. After 72 hours of treatment, cell proliferation was assessed using the the BrdU Cell Proliferation ELISA kit (Roche, Basel, Switzerland) according to the manufacturer’s protocol. Optical density was measured at 450 nm using a microplate reader (Bio-Rad, Hercules, CA, USA).

### Cell apoptosis analysis

Apoptosis was evaluated using the Annexin-V-FLUOS Staining Kit (cat. 11 858 777 001; Roche) in accordance with the manufacturer’s instructions. FLS (2 × 10^5^ cells/well) were plated in six-well plates. Following treatment, suspended cells were collected, and adherent cells were detached by Accutase® solution (Sigma-Aldrich). Both cell populations were washed with cold PBS and resuspended in 100 µl Annexin-V-FLUOS labeling solution (containing 2 µl Annexin-V-FLUOS reagent and 2 µl Propidium Iodide). After incubation in the dark for 15 minutes at room temperature, cells were analyzed by flow cytometry (FACSVerse, BD Biosciences).

### Mice

Female SKG mice, a murine model of rheumatoid arthritis (RA), were purchased from CLEA Japan, Inc. The mice were housed at the Institute of Laboratory Animals, Kobe University. All animal studies were conducted in accordance with the Institutional Animal Care and Use Committee guidelines of Kobe University and approved by the same committee (approval number: P191009). Mice were sacrificed by subcutaneous injection of pentobarbital, following institutional ethical guidelines.

### Arthritis induction and evaluation

Arthritis was induced in SKG mice by intraperitoneal injection of 2 mg Zymosan A (ZyA; AlfaAesar, Lancashire, UK) suspended in 500 µl saline at eight weeks of age. Arthritis onset occurred 15–27 days post-injection. The arthritis score was determined using a previously established scoring system [22,23]:

・0 = no joint swelling

・0.1 = swelling of a single digit joint

・0.5 = mild swelling of the wrist or ankle

・1.0 = severe swelling of the wrist or ankle

Scores were summed for each mouse, with a maximum possible score of 5.8.

### Treatment of SKG mice with CQ and C968

Chloroquine (CQ; Sigma-Aldrich, St. Louis, MO, USA) was dissolved in saline, while 500 µg of C968 (Calbiochem; La Jolla, CA, USA) was dissolved in 50 µl dimethyl sulfoxide (DMSO; Tocris Bioscience, Bristol, UK). Once the average arthritis score reached 0.25 (27 days post-ZyA injection), mice were randomly assigned to four treatment groups and received subcutaneous injections as follows:

1. Vehicle: saline daily + DMSO three times per week
2. CQ: CQ (25 mg/kg, daily) + vehicle (DMSO three times per week)
3. C968: C968 (25 mg/kg, three times per week) + vehicle (saline daily)
4. CQ + C968: CQ (25 mg/kg, daily) + C968 (25 mg/kg, three times per week)

All treatments were administered via subcutaneous injection for 28 days.

### Statistical analysis

Data are presented as mean ± SEM. Statistical analyses were performed using an unpaired Student’s t-test and one-way ANOVA followed by post-hoc analysis. A p-value < 0.05 was considered statistically significant.

## Results

### C968 induces autophagy in RA-FLS

To investigate the effect of C968, a GLS1 inhibitor, on autophagy in RA-FLS, we evaluated the expression of autophagy-related genes, ATG5 and LC3B, using real-time PCR following treatment with C968 (0, 20 and 30 μM). C968 upregulated ATG5 and LC3B expression in a dose-dependent manner (Fig 1A). Next, we examined LC3-II protein levels in RA-FLS after treatment with 20 μM C968 or 10 μM chloroquine (CQ) via western blotting. LC3-II is present on autophagosome membranes and its amount reflects the number of autophagosomes. We observed a progressive increase in LC3-II levels at 24 and 48 hours post-C968 treatment (Fig 1B). Additionally, CQ, a lysosomal inhibitor that prevents autophagosome-lysosome fusion, increased LC3-II levels at 48 hours post-treatment.

**Fig 1.**
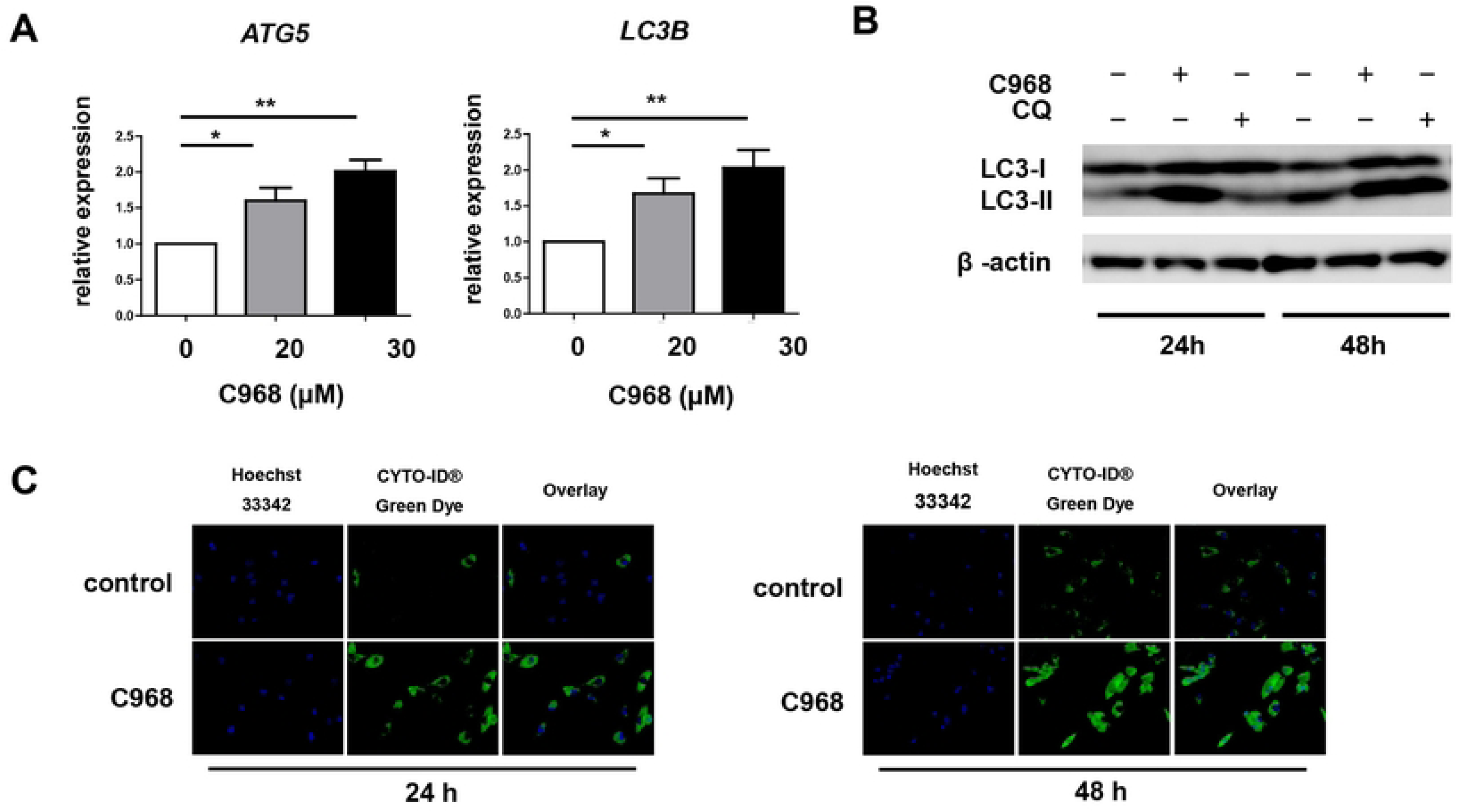
C968 induces autophagy in rheumatoid arthritis fibroblast-like synoviocytes (RA-FLS). (A) RA-FLS were incubated with different concentrations (0, 20, and 30 μM) of C968 for 48 h. mRNA expression levels of ATG5 and LC3B were quantified by real-time PCR and normalized to GAPDH levels. Bar graphs indicate the mean ± SEM. ATG5 data are from four independent experiments, and LC3B data are from six independent experiments. *p < 0.05; **p < 0.01 (one-way ANOVA followed by Dunnett’s multiple comparison test). (B) RA-FLS were cultured with 20 μM C968 or 10 μM CQ for 24 and 48 h. LC3 and β-actin levels were determined by western blot analysis. (C) RA-FLS were cultured with or without 20 μM C968 for 24 and 48 h. FLS were then stained with Cyto-ID® Green Dye and analyzed using confocal fluorescence microscopy.

To further confirm C968-induced autophagy, we performed immunofluorescence staining to visualize autophagosome formation in RA-FLS treated with 20 μM C968. C968-treated FLS exhibited a pronounced autophagosome-specific green fluorescence signal at both 24 and 48 hours, compared with untreated cells (Fig 1C).

These findings confirm that C968 induces autophagy in RA-FLS.

### Inhibition of C968-induced autophagy suppresses cell growth and promotes apoptosis in RA-FLS

Autophagy is known to play a cytoprotective role in various diseases, particularly in cancer, where its inhibition has been shown to enhance therapy-induced cell death. To investigate whether inhibiting C968-induced autophagy affects RA-FLS proliferation and apoptosis, we examined the combined effects of C968 and CQ, an autophagy inhibitor, on cell growth and apoptosis.

As shown in Fig 2A, C968 treatment alone reduced RA-FLS proliferation, and co-treatment with C968 and CQ further suppressed cell proliferation. Moreover, the combination of C968 and CQ significantly increased apoptosis, whereas neither C968 nor CQ monotherapy had a comparable effect (Figs 2B and 2C).

**Fig 2.**
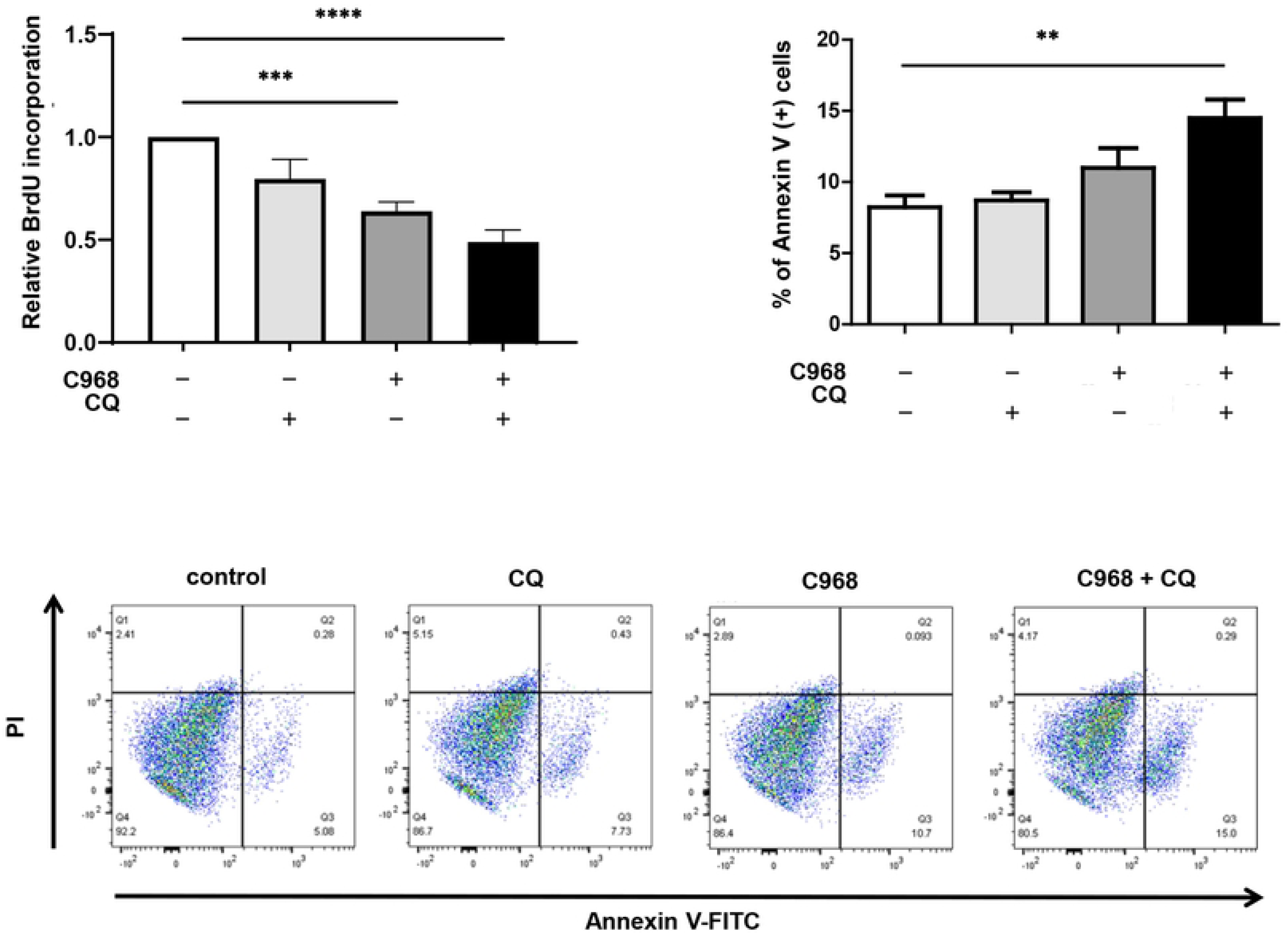
Inhibiting C968-induced autophagy suppresses cell growth and promotes apoptosis in RA-FLS. (A) RA-FLS were treated with 10 μM C968 in the presence or absence of 30 μM CQ for 72 h. Cell proliferation was measured by BrdU assay. Each experiment was performed in quintuplicate. Data are presented as the mean ± SEM. *p < 0.05; ***p < 0.001 (one-way ANOVA followed by Dunnett’s multiple comparison test). (B) RA-FLS were treated with 30 μM C968 with or without 20 μM CQ for 24 h. Apoptosis was determined using Annexin-V-FLUOS and propidium iodide (PI) staining by flow cytometry. Data are expressed as the percentage of Annexin-V-positive cells. Bars represent the mean ± SEM of five independent experiments. **p < 0.01 (one-way ANOVA followed by Dunnett’s multiple comparison test). (C) Representative flow cytometry images corresponding to Fig 2B.

These results suggest that inhibiting C968-induced autophagy more effectively suppresses cell growth and promotes apoptosis in RA-FLS.

### Dual suppression of autophagy and glutaminolysis ameliorates arthritis in SKG mice

Finally, we investigated the in vivo effects of dual suppression of autophagy and glutamine metabolism. SKG mice with ZyA-induced arthritis received subcutaneous injections of CQ, C968, or a combination of both for 28 days.

The combination of C968 and CQ significantly reduced arthritis severity in SKG mice (Fig 3), whereas neither C968 nor CQ monotherapy resulted in a significant improvement (S1 Fig).

**Fig 3.**
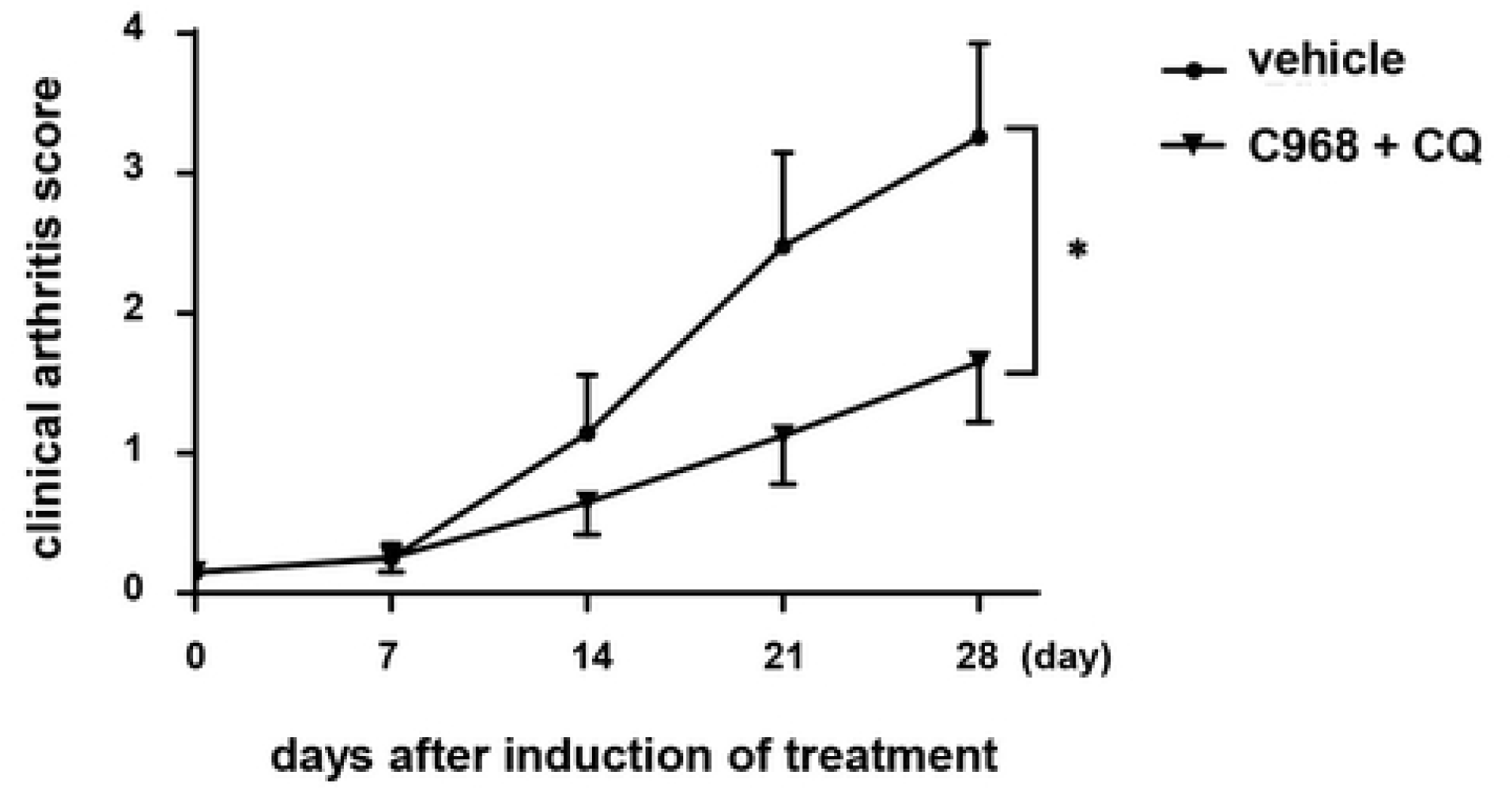
Suppressing autophagy and glutaminolysis ameliorates arthritis in SKG mice. SKG mice with ZyA-induced arthritis were subcutaneously injected with vehicle (n = 10) or CQ in combination with C968 (n = 15). Clinical arthritis scores were recorded for up to 28 days following drug administration. *p < 0.05 on day 28 (Student’s t-test).

These findings indicate that dual inhibition of autophagy and glutaminolysis may represent a promising therapeutic approach for autoimmune arthritis.

## Discussion

RA is a multifactorial and heterogeneous disease, and significant unmet clinical needs persist [6]. Therefore, the development of novel and effective therapeutic approaches remains an urgent priority. The antimalarial drug CQ and its derivative hydroxychloroquine (HCQ)—which share similar pharmacokinetics and mechanisms of action—have long been used to treat RA and other autoimmune diseases [7]. However, CQ or HCQ monotherapy has shown only limited efficacy in RA, necessitating combination therapy [8].

Recent research has focused on the anti-autophagic effects of CQ and HCQ. In certain cancers, chemotherapy-induced autophagy contributes to drug resistance, and autophagy inhibition via CQ or HCQ has been shown to enhance anticancer therapeutic efficacy [12].

Here, we demonstrate for the first time the superior efficacy of the CQ–C968 combination in suppressing RA-FLS proliferation and ameliorating arthritis in SKG mice, compared with monotherapy alone. Our study revealed that glutaminolysis inhibition by C968 enhanced autophagy in RA-FLS. Although the precise mechanistic link between glutamine metabolism and autophagy remains incompletely understood, glutaminolysis has been proposed to suppress autophagy via mTORC1 activation or by counteracting ROS production [20]. In cancer models, inhibition of glutaminolysis or its downstream target, mTORC1, has been shown to induce compensatory autophagy, limiting its therapeutic effect, and demonstrating synergistic efficacy when combined with autophagy inhibition [24–26]. Consistent with these findings, our study showed that co-treatment with C968 and CQ significantly reduced RA-FLS proliferation, enhanced apoptosis, and mitigated arthritis in SKG mice, suggesting that CQ-mediated suppression of C968-induced autophagy enhances therapeutic efficacy (Fig 4).

**Fig 4.**
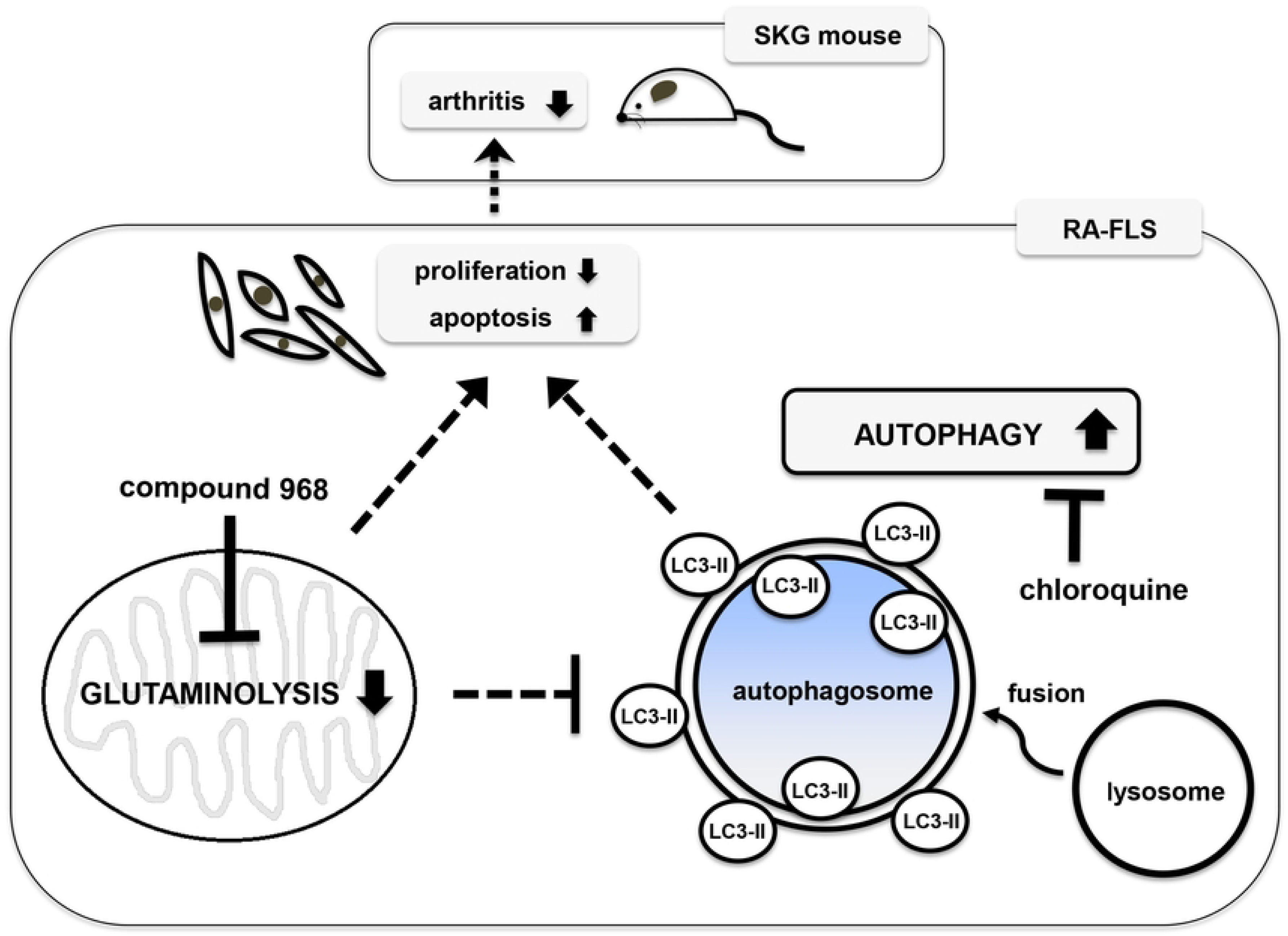
Major mechanisms of C968 and chloroquine treatment in RA-FLS.

Inhibition of glutaminolysis by C968 reduces cell proliferation and promotes apoptosis in RA-FLS. However, glutaminolysis inhibition induces autophagy, which may support the proliferative and anti-apoptotic characteristics of RA-FLS. Combined treatment with C968 and CQ, an autophagy inhibitor, enhances the therapeutic effect of C968 on RA-FLS and alleviates arthritis in SKG mice.

Although our previous study demonstrated that C968 attenuates arthritis in SKG mice [19], C968 monotherapy failed to show a significant effect in the present study. This discrepancy may be attributed to differences in administration routes. In our previous study, C968 was administered via intraperitoneal injection, whereas in this study, subcutaneous injection was used, as it is more relevant for potential clinical applications. Given that C968 is highly hydrophobic, containing five benzene rings and lacking hydroxyl groups, its absorption and tissue distribution may be lower following subcutaneous administration compared to intraperitoneal injection. This pharmacokinetic difference may explain the lack of efficacy observed with C968 monotherapy in this study, although pharmacokinetic data on C968 in relation to different administration routes remain unavailable.

This study has some limitations. While both glutaminolysis and autophagy have been implicated in T cell activity [27–31], we did not assess the impact of C968 or CQ on immune cell function. Given that glutaminolysis is a key metabolic pathway for T cell activation [27] and that autophagy plays a crucial role in T cell proliferation and survival [28–31], the observed therapeutic effects of C968 and CQ in SKG mice may not be solely attributed to RA-FLS regulation but also to immune cell modulation. Furthermore, CQ and HCQ possess diverse immunomodulatory properties beyond autophagy inhibition, including Toll-like receptor inhibition [32], reduction of proinflammatory cytokine secretion [33], downregulation of TNF receptors [34], and impairment of antigen presentation [35, 36]. However, this study primarily focused on the autophagy-inhibitory effects of CQ. Future studies are warranted to comprehensively elucidate the broader immunomodulatory effects of these compounds.

Although CQ’s additional effect on C968 treatment may not be solely attributed to autophagy inhibition, our in vitro and in vivo findings strongly suggest that simultaneous suppression of C968-induced autophagy is essential to potentiate the therapeutic effects of C968 on arthritis.

## Conclusion

In this study, we demonstrated that the GLS1 inhibitor C968 enhances autophagy in RA-FLS. Furthermore, we showed that co-treatment with C968 and CQ synergistically suppressed RA-FLS proliferation, enhanced apoptosis, and significantly ameliorated arthritis in SKG mice.

These findings suggest that dual inhibition of glutaminolysis and autophagy represents a promising therapeutic strategy for RA.

## Acknowledgments

We sincerely thank Shino Natsui for her invaluable technical support. We also express our gratitude to Editage for their assistance with English language editing.

---

**Fig S1.**
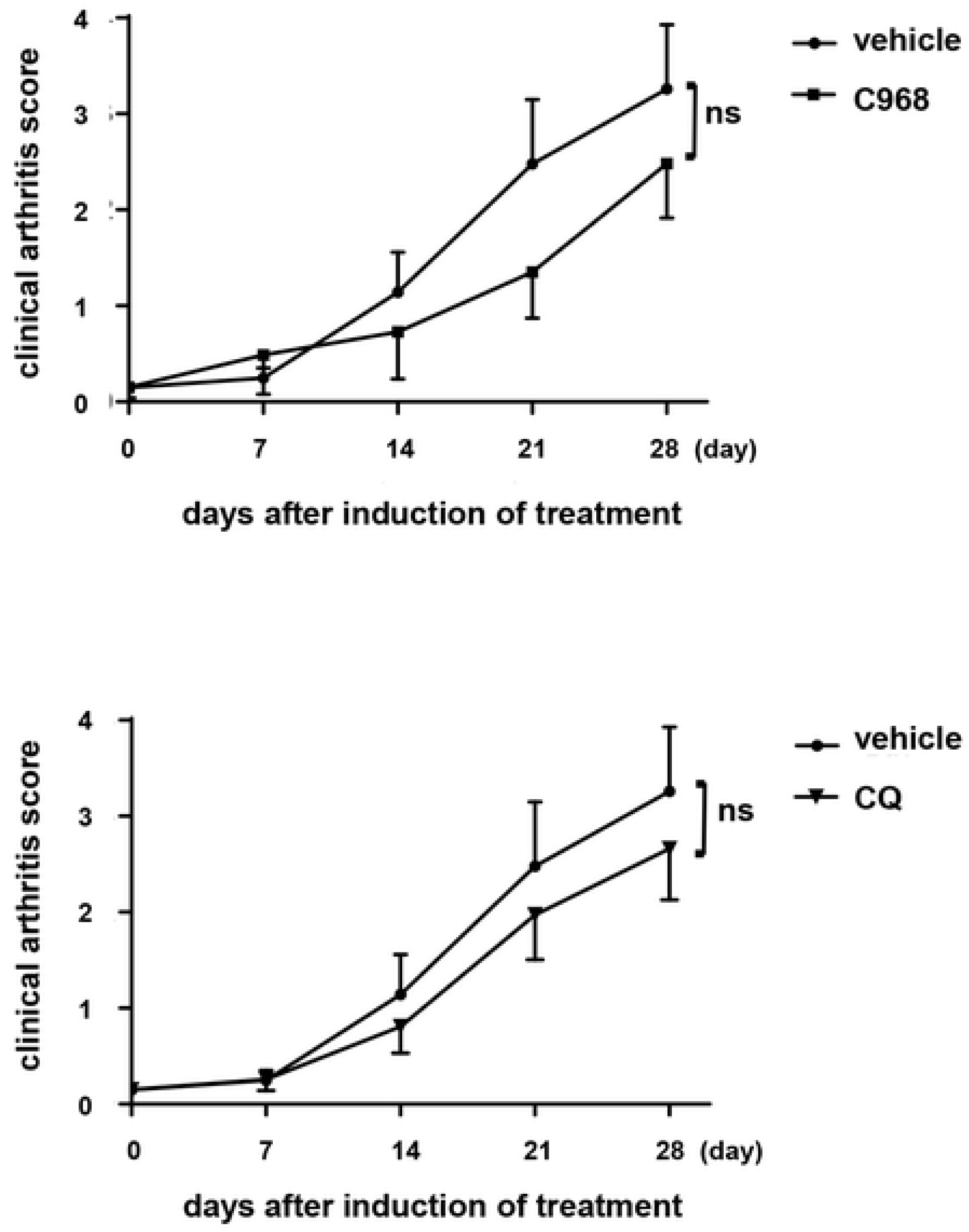
Neither C968 nor CQ monotherapy significantly attenuates arthritis severity in SKG mice. SKG mice with ZyA-induced arthritis were given subcutaneous injections of vehicle (n = 10), C968 (n = 14), or CQ (n = 10). Clinical arthritis scores were recorded for up to 28 days following the initiation of drug administration.

## Notes

### Competing Interest Statement

The authors have declared no competing interest.

